# Characterizing Mineral Ellipsoids in New Bone Formation at the Interface of Ti6Al4V Porous Implants

**DOI:** 10.1101/2023.01.20.524810

**Authors:** Joseph Deering, Jianyu Chen, Dalia Mahmoud, Tengteng Tang, Yujing Lin, Qiyin Fang, Gregory R. Wohl, Mohamed Elbestawi, Kathryn Grandfield

## Abstract

The hierarchy of newly formed bone contains elements of disorder within an ordered multiscale structure, spanning from the macroscale to below the nanoscale. With mineralized structures presenting in the shape of ellipsoids in mature and mineralizing tissue, this work characterizes the heterogeneity in mineral ellipsoid packing at the interface of porous titanium implants. The characterization of mineral at the bone-implant interface offers insight into the osseointegration of titanium implants and the mechanical properties of the interfacial bone tissue. Using scanning transmission electron microscopy and plasma focused ion beam - scanning electron microscopy, mineral ellipsoids are characterized at the implant interface in both 2D and 3D. Heterogeneous in their size and shape within the newly formed bone tissue, ellipsoids are observed with alternating orientations corresponding to unique lamellar packets within 2-3 μm of the titanium implant interface – although this motif is not universal, and a mineral-dense zone can also appear at the implant interface. Short-order ellipsoid orientation shifts are also present in the 3D probe of the implant interface, where an approximate 90° misorientation angle between neighbouring packets of mineral ellipsoid resolves with increasing distance from the titanium, possibly providing a strengthening mechanism to prevent crack propagation in the peri-implant bone.

**Graphical Abstract:** 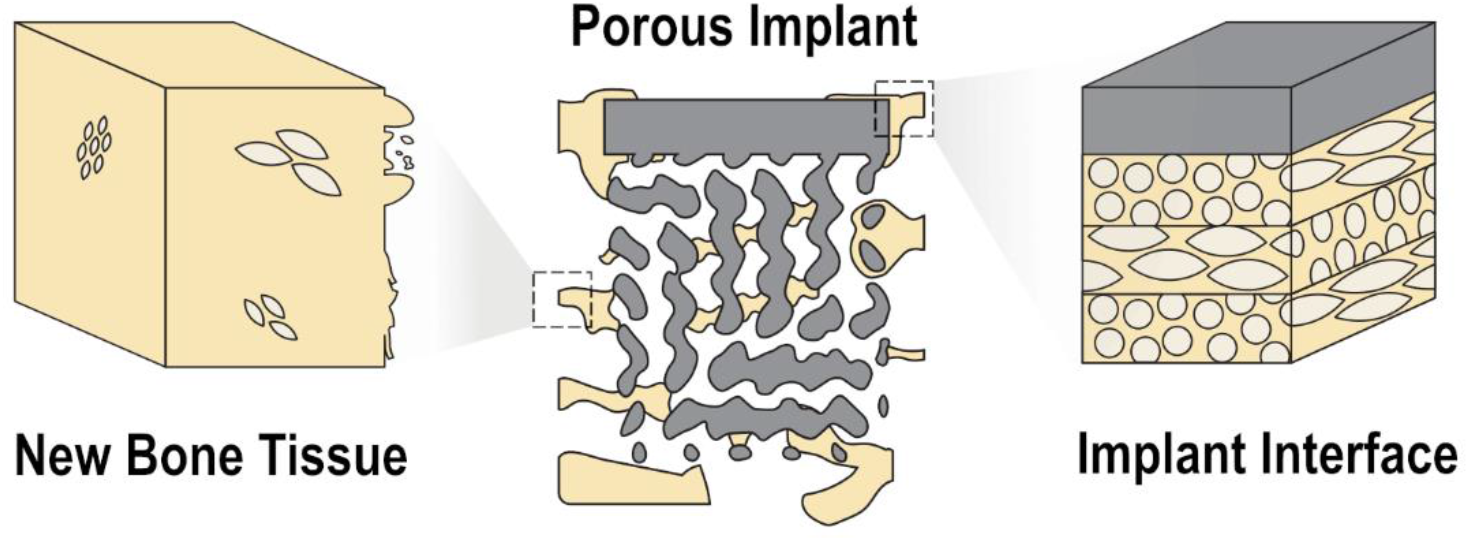

## 1 Introduction

The natural structure of mineralized bone tissue spans the continuum between the macroscale and nanoscale, encompassing both organic and inorganic components to achieve robust mechanical properties. At smaller length scales, osteoblast-secreted type I collagen constitutes much of the organic assembly and provides a template for mineral ingress into the tissue ^[1]^. As the tissue begins to mineralize, an ellipsoidal mineral packing arrangement is theorized to occur according to a stencilling principle based on inhibition of inhibitors ^[2]^, with resulting mineral clusters sized at roughly 0.4-1.5 μm along their major and minor axes ^[3]^. When viewed in orthogonal directions with two-dimensional imaging such as transmission electron microscopy (TEM), these ellipsoids take on either an elliptical motif or a circular rosette motif. Based on large-scale 3D imaging, these mineral clusters appear to be ubiquitous throughout bone tissue (as noted in human femora ^[4]^ and murine tibiae ^[5]^) and have been noted adjacent to implants of titanium, bioactive glass, and others in either of the elliptical or rosette motifs ^[3]^. Yet, little is known about their shape, origin, or evolution in new bone tissue or their 3D organization at the interface of metallic biomaterials despite a wealth of literature about smaller features such as collagen orientation ^[6,7]^.

Additive manufacturing offers a means to produce implants with engineered porosity in the microscale range ^[8]^, with porous titanium and tantalum implants exhibiting mechanical properties similar to cortical bone ^[9]^ and improved secondary stability in biomechanical evaluation ^[10]^. The low mechanical stiffness ^[11,12]^ imparted by these implants offers one method to mitigate long-term bone resorption associated with stress-shielding ^[13]^ and the corresponding increase in surface area can create additional regions of bone-to-implant contact ^[14]^. Understanding bone mineral apposition and morphological transformations at implant interfaces and into these pores may therefore have major implications for early osseointegration, implant stability, and longevity of implants in the body.

Prior to the discovery of mineral ellipsoids, osseointegration at implant interfaces was identified by key markers and used techniques other than electron microscopy ^[15,16]^. At the microscale, observations of cellular response ^[17]^ or woven-to-lamellar transition ^[18,19]^ are often used to qualitatively and quantitatively assess osseointegration. At the nanoscale, higher-resolution studies have shown the accumulation of non-collagenous proteins near the implant interface ^[20,21]^ or that of a calcified cement line ^[22]^, and generally denoted the orientation of collagen parallel to the implant surface ^[23,24]^. While all of these observations give a strong understanding of bone growth near implants, a lack of widespread 3D imaging may be at the heart of why mineral ellipsoids have not been studied in detail at implant interfaces, with few examples of osseointegration studies using FIB-SEM over small volumes ^[25]^, and electron tomography with an even smaller field of view^[26–30]^. Moreover, microscale pores present a unique challenge for characterization, with a simultaneous need for high-resolution and large-volume imaging. Plasma focused ion beam tomography provides a window to examine tissue apposition in the pore network over time.

Here, we report a high resolution and 3D study of the osseointegration of titanium implants with a graded pore size in rabbits at 4 and 12 weeks. We utilize scanning transmission electron microscopy and plasma focused ion beam to provide an innovative approach for assessing ellipsoidal mineral organization at implant interfaces – achieving both high-resolution and 3D structure - and offering insight into potential consequences with respect to the mechanical properties of interfacial bone tissue.

## 2 Methods

### 2.1 Scaffold Design and Implantation

Porous implant scaffolds were designed and implanted according to the specifications described elsewhere ^[31]^. Briefly, implants were designed with a total height of 7 mm and total diameter of 6 mm. The porous midsection was designed to contain a gyroid unit cell, with radially varying pore size from *ϕ* = 600 μm at the scaffold exterior to *ϕ* = 300 μm at the midpoint of the scaffold. Powder bed fusion of Ti6Al4V powder feedstock was used to create the implants (Renishaw, Kitchener, ON, Canada).

Under general anesthesia, implants were placed bilaterally into the tibial epiphyses of 6-month old male New Zealand white rabbits (2.5-3 kg) with approval from the Animal Care Committee of Sun Yat-sen University (Approval No. SYSU-IACUC-2020-000198). Antibiotics were administered for three days following animal surgery. One implant was retrieved after 4 and a second after 12 wk to prepare for electron microscopic analysis.

### 2.2 Sample Preparation

Soft tissue attached to the sample was carefully removed with a scalpel, and full tibiae were defatted after implant retrieval using immersion in 100% acetone for 12-24 hr. Samples were then fixed in 10% neutral buffered formalin (Sigma Aldrich, St. Louis, MO, USA) on a rocking table for 24 hr before washing with deionized water three times (4 hr, 4 hr, and 12 hr each) to remove fixative and subsequent sectioning to the implant area using a fret saw. Following sectioning, serial dehydration was conducted for each specimen using 48 hr intervals at 50%, 70%, 70%, 80%, 90%, 95%, 95%, 100%, 100% ethanol concentrations under vacuum (−80 kPag). Afterwards, Embed 812 resin (Electron Microscopy Sciences, Hatfield, PA, USA) was gradually infiltrated into the specimens in 48 hr intervals using ratios of 1:3, 1:1, and 3:1 in acetone as a solvent. Specimens were transferred to 100% Embed 812 resin in embedding molds and cured at 60 °C under vacuum (−90 kPag) for 48 hr. Embedded implants were sectioned along the axial plane at their approximate midpoint using a slow-speed diamond saw (IsoMet Low Speed Saw, Buehler, Lake Bluff, IL, USA) to reveal the to achieve a flat surface. Finally, the specimens were mounted to aluminium stubs using silver paint (Ted Pella Inc., Redding, CA, USA) and sputter coated with gold (~ 20 nm thick) to increase conductivity. A scanning electron microscope (JEOL 6610LV, Tokyo, Japan) was used to examine the specimen surface with an acceleration voltage of 20 kV, working distance of 10 mm, and spot size of 60. Areas where the implant interfaces with bone tissue were selected for further analysis.

### 2.3 Scanning Transmission Electron Microscopy

A site of bone-implant contact was identified using SEM near the crown of the implant following twelve weeks of implantation (**Figure S1**). To prepare a specimen for scanning transmission electron microscopy (STEM), a FIB milling and lift-out technique was used, following a procedure described elsewhere ^[32]^. Briefly, a Zeiss NVision 40 (Carl Zeiss, Oberkochen, Germany) was used to prepare a ~ 100 nm thick specimen at the interface of the implant. A 4 μm thick layer of tungsten was initially deposited over the region of interest followed by coarse and fine trench milling on all sides at 30 kV with currents ranging from 6.5 to 45 nA. The specimen, containing bone flanked on either side by titanium implant, was then lifted out of the bulk sample and attached to a copper grid, where electron transparency (70-120 nm thick) was achieved using a series of decreasing ion beam currents from 300 pA to 40 pA at 30 kV. The final ion polishing of the thinned specimen was conducted at an acceleration voltage of 5 kV to remove surface unevenness and redeposition.

STEM examination was conducted in a Talos 200X (ThermoFisher Scientific, Waltham, MA, USA) using a high-angle annular dark field (HAADF) detector and an accelerating voltage of 200 kV with eightfold frame averaging. Electron energy loss spectroscopy (EELS) was conducted using a dispersion of 0.3 eV/channel, acquisition time of 0.01s, and pixel size of 7 nm on a Continuum S imaging filter (Gatan). EELS elemental maps for carbon, calcium, and titanium were generated in Digital Micrograph software (Gatan Inc., Pleasanton, CA, USA).

### 2.4 Plasma Focused Ion Beam-Scanning Electron Microscopy

The implants retrieved after four weeks and twelve weeks were polished further with silicon carbide paper to expose a suitable site for plasma focused ion beam - scanning electron microscopy (PFIB-SEM) tomography. The two specimens (4 wk and 12 wk) were loaded into a Helios G4 UXE DualBeam (ThermoFisher Scientific) microscope with a xenon plasma source for serial sectioning. Sites were selected according to the schematic in **Figure 1**, sampling both cortical and trabecular bone-implant interfaces at a radial depth of roughly 150 μm into the implant interior (**Figure S1 and S2**).

**Figure 1.**
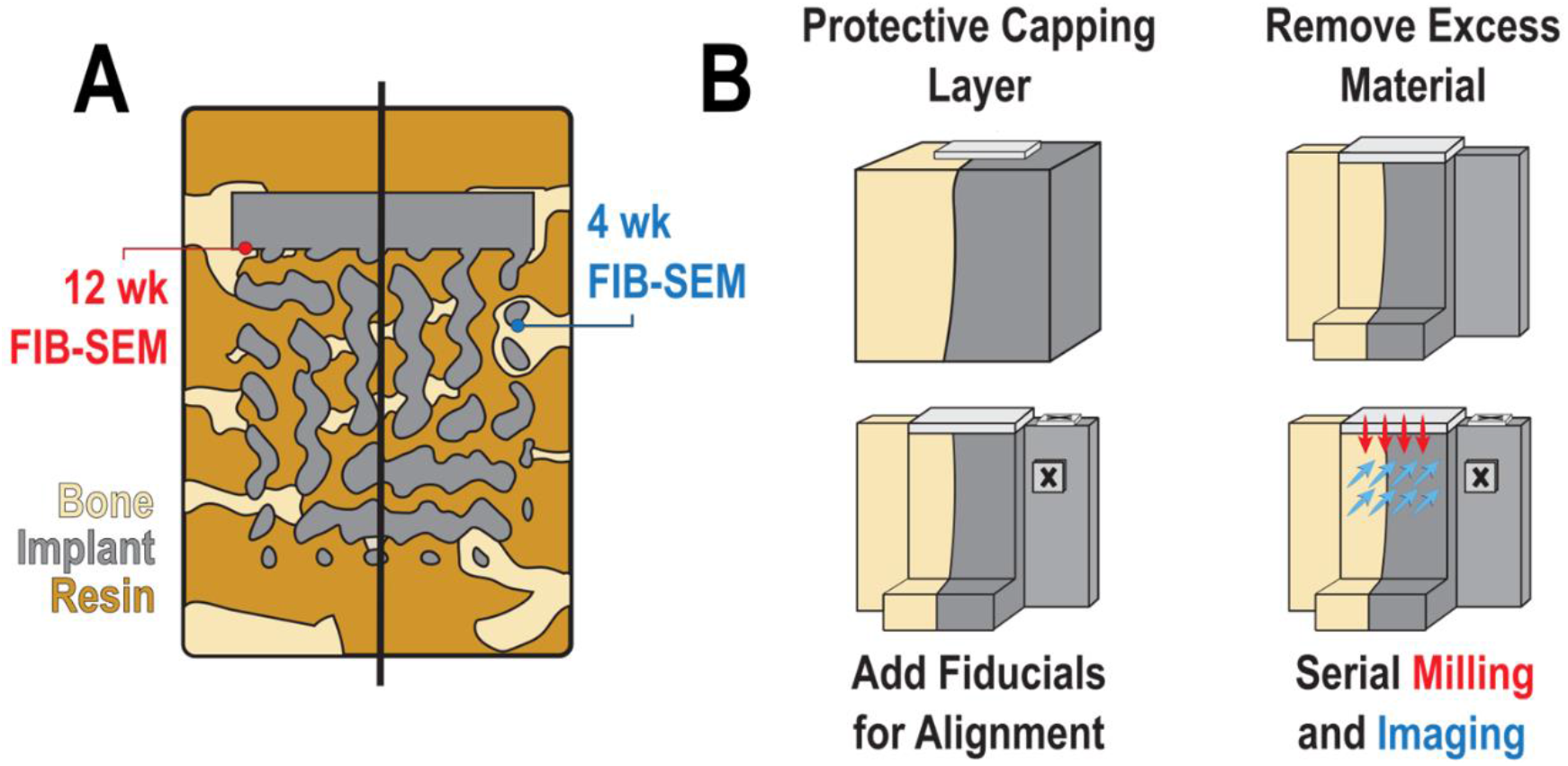
Workflow of PFIB-SEM tomography at the bone-implant interface. (A) A schematic showing regions of interest extracted at near-surface implant pores in the cortical region after 12 wk and at regions of trabecular ingrowth after 4 wk of implantation. (B) Serial sectioning of PFIB-SEM takes place after the region of interest is carefully prepared using sequential deposition of a protective capping layer, removal of excess material, and addition of fiducials for ion beam tracking and postprocess image registration.

Briefly, a protective carbon layer (10-13 μm thick) was deposited on the specimen surface to minimize ion beam damage and curtaining artifacts. Excess material was subsequently removed around the volume using beam currents of 15-200 nA at 30 kV to create trenches, and fiducials were added to the top surface and the block-face onto small pads of tungsten deposition. Auto Slice and View (ThermoFisher Scientific) was used to automate the sequential milling and imaging processes for tomography acquisition. Specifically, milling was performed at 30 kV and 1-4 nA with the ion beam using a 4° rocking angle and slice thickness of ~30 nm. Images in the series were acquired with a 1.2 kV, 400 pA electron beam in immersion mode using a 500 ns dwell time and pixel size of 14.5 nm × 14.5 nm, with fourfold line integration and averaging across two frames on the through-the-lens detector (6144 × 4096 pixels).

### 2.5 PFIB-SEM Data Processing

All image processing and segmentation was conducted in Dragonfly 2022.1 (Object Research Systems, Montreal, Quebec, Canada). PFIB-SEM tomography datasets were aligned using mutual info from the block-face fiducial to translate slices with a 5.0% initial translation step in the imaging plane and smallest translation step of 0.05%. Datasets were subjected to a 3D Gaussian blur (kernel size = 5, σ = 1.0) without contrast correction before further processing.

#### 2.5.1 Segmentation and Analysis of Bone Mineral Ellipsoids

Bone tissue adjacent to the titanium implant in the 4 wk PFIB-SEM dataset was extracted using grayscale-based segmentation, with a threshold value of 155 or greater in the 8-bit images. Clusters or aggregated clusters of mineral near the bulk of the bone tissue were segmented using 6-connected component labelling to isolate their boundary from any adjacent mineralized entities. Mineral clusters were delineated using a size threshold of 500 voxels or greater..

#### 2.5.2 Segmentation of Bone Lacunocanalicular Network

The lacunocanalicular network (LCN) within the mineralized bone of the 4 wk PFIB-SEM datasets was segmented by further Gaussian smoothing (2D, kernel size = 15, σ = 4.0) and using application of grayscale-based thresholding tool (0-150) to extract the LCN and features with similar grayscale value in the implant retrieved after 4 wk. Bone matrix (along with other high-contrast regions) was extracted in this PFIB-SEM dataset using a grayscale window of 150-256, with a subsequent fill operation in the Z-direction to infill the darker LCN components (including regions of charging). A Boolean intersection was taken between the matrix/titanium/LCN segmentation and resin/LCN segmentations for final extraction of the LCN components. The LCN in the 12 wk PFIB-SEM dataset was segmented in a similar grayscale-based manner across four partitions of the dataset to account for contrast changes arising from microscope instability.

## 3 Results and Discussion

### 3.1 Chemistry of the Bone-Implant Interface

Understanding the structural and compositional interactions between bone and an adjacent implant surface can help provide insight into the osseointegration and mechanical stability of the implant. To this end, scanning transmission electron microscopy (STEM) and electron energy loss spectroscopy (EELS) can provide nanometer scale spatial and chemical information about the bone-implant interface in 2D. A representative high-angle annular dark-field (HAADF) STEM image at the interface of a porous titanium implant retrieved after 12 wk is shown in **Figure 2A**, where electron dense regions resembling the shape of ellipsoids were observed near the implant interface but not in the most interfacial region of bone tissue. These dense mineralized regions of the ellipsoid interior were separated by a less mineralized fibrillar matrix.

**Figure 2.**
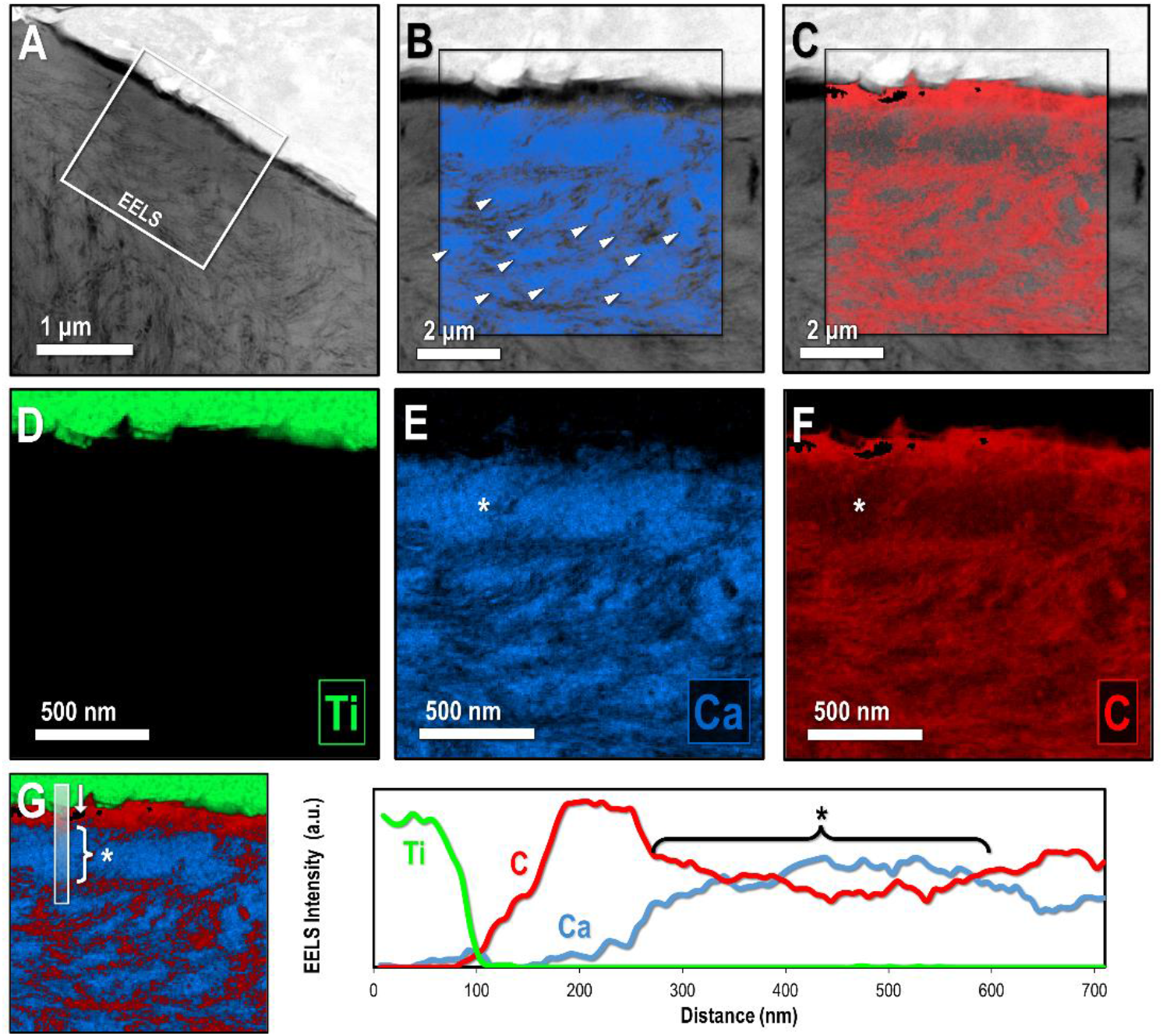
STEM-EELS of de novo bone formation at the implant interface. (A) Site overview of bone-implant near cortical interaction with the crown of the implant after 12 wk. (B-C) Calcium and carbon EELS maps overlaid with the STEM image at the implant interface. Arrowheads represent individual mineral ellipsoids. (D-F) Titanium, calcium, and carbon EELS maps at the implant interface. Carbon-rich areas tend to overlap with calcium-deficient regions. A mineral-dense zone (*) is present in the tissue at the interface of the titanium. (G) EELS intensity profiles across the mineral-dense zone. Marginal carbon depletion occurs in this mineral zone in addition to slight calcium elevation.

Corresponding EELS maps of calcium (**Figure 2B** and **E**), carbon (**Figure 2C** and **F**), and titanium (**Figure 2D**) provide information on chemical gradients across the interface. Within the first 150 nm of the titanium interface, there is an increased level of carbon content, which is mostly likely attributed to the resin shrinkage during tissue preparation. Nonetheless, the chemical map of the bone-implant interface shows distinct carbon-rich and calcium-rich regions near the implant. In the 300 nm beyond this preparation artifact, there is first a zone of higher mineral density with slightly elevated calcium and slightly depleted carbon (**Figure 2G**). Such observations agree with previous reports where discrete regions with a seemingly afibrillar appearance are seen within the first few nanometres of the implant interface, manifesting as a cement line ^[33]^, but not readily displaying the mineral ellipsoid motif. An afibrillar appearance at the interface was proposed to attribute to either the adhesion of early non-collagenous protein on the implant surface to promote mineralization with the assembly of a collagenous layer shortly following ^[34]^, but the deposition of a dominant collagenous zone has also been seen in the direct vicinity of the implant with only a small fraction of granular or afibrillar material ^[24]^. In either case, the structure and packing density of mineral at the bone-implant interface can give prospective information about the assembly and transformation of bone-implant contact during osseointegration. It is also important to note that this mineral-dense band adjacent to the implant does not span the entire titanium surface but appears sporadically across the bone-implant interface and has not been seen next to modified implant surfaces ^[35]^.

Further from the implant interface, the fibrillar appearance of carbon-rich zones loosely delineate the boundary between mineral ellipsoids – where the ellipsoid interior appears marginally higher in calcium content than at its boundary. These distinct collagen fibril bundles can be observed outside of the mineral-dense band, where mineral ellipsoids appear more loosely packed. While the mineral-dense band likely still contains collagen fibrils in some quantity, the size distribution or high packing density of mineral ellipsoids within this band could mask the collagen in HAADF-STEM imaging but it is also important to consider that this phenomenon could arise due to variable thickness in the thinned region of interest.

### 3.2 Evidence of Formative Processes at the Bone-Implant Interface

PFIB-SEM tomography of the bone-implant interface provides a nanometer resolution 3D probe that can span microscale volumes of biological tissues, with faster milling rates using the xenon beam than conventional gallium-based ion beams [36]. 3D volumes of this size (56.2 μm x 57.0 μm x 14.0 μm at 4 wk and 68.9 μm x 59.1 μm x 11.4 μm at 12 wk) provide useful insight into components of mineralized tissue – spanning regions of the implant itself (bright regions), the mineralizing bone tissue (intermediate regions), and carbon-rich or porous entities (dark regions) (**Figure 3**).

**Figure 3.**
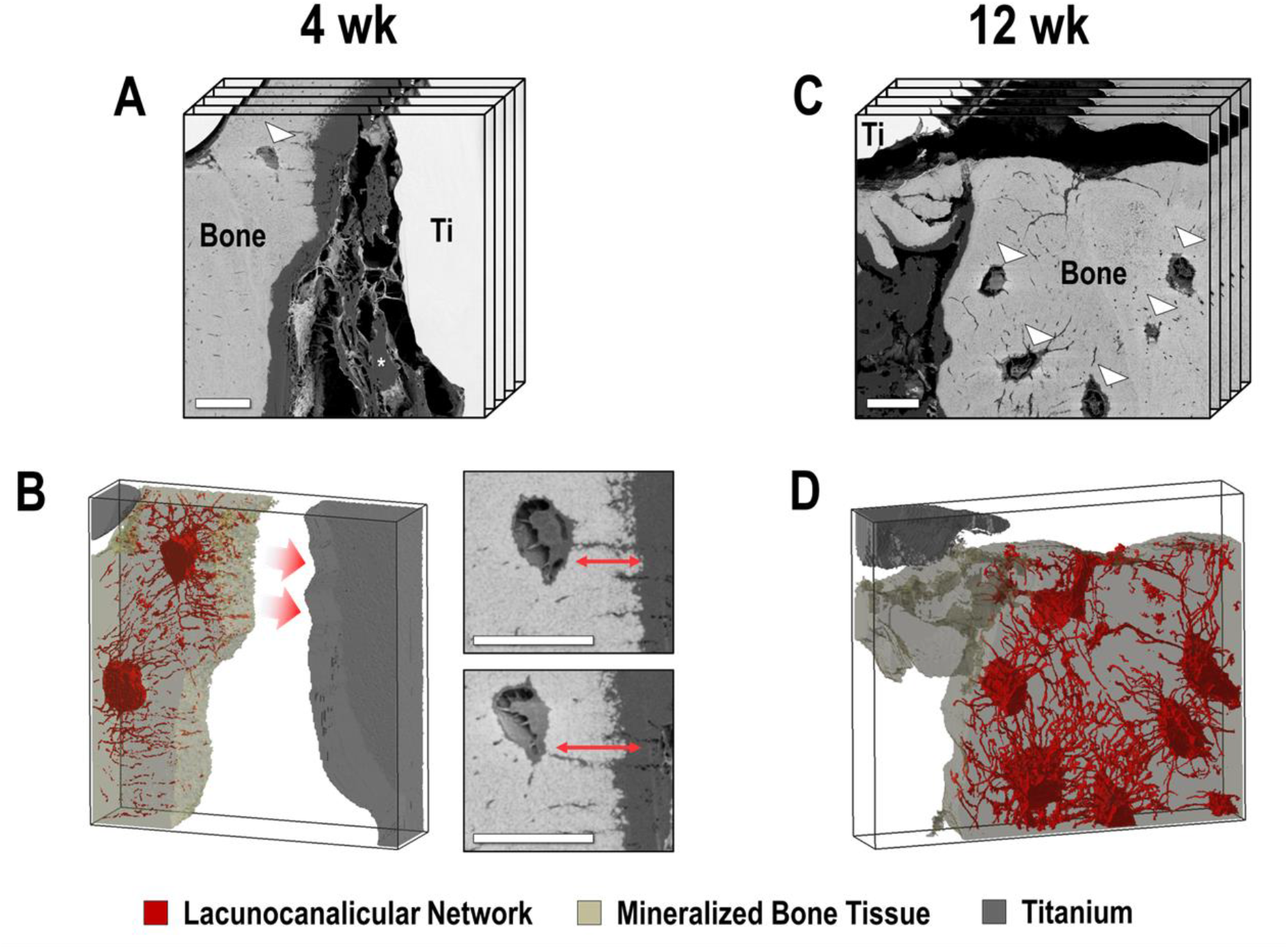
Representative slices from PFIB-SEM tomography and osteocyte reconstructions. (A) 3D representation of the PFIB-SEM image stack of the implant interface retrieved after 4 wk. Brightest regions are indicative of titanium, dark regions are indicative of carbon- or resin-rich material, and intermediate grayscale regions are mineralized bone tissue. Osteocyte lacunae are denoted with arrowheads and bone-implant contact is evident in the upper left quadrant. (B) Segmentation of the bone matrix, implant volume, and LCN after 4 wk shows regions where canaliculi predominantly extend towards the edge of the mineralized bone, indicating active growth towards the titanium implant. (C) 3D representation of the PFIB-SEM image stack at the implant interface acquired 12 wk after implantation. (D. After 12 wk, the dense cell network is distributed through the bone, with no clear regions of preferential orientation. Scale bars: 10 μm.

PFIB-SEM reconstructions of the bone-implant interface after 4 wk and 12 wk of implantation in leporine tibia is shown in **Figure 3A** and **Figure 3C**, respectively. Corresponding segmentation of each component (mineralized bone matrix, lacunocanalicular network, and titanium) is shown in **Figure 3B** and **Figure 3D**. After 4 wk of implantation, bone in the shallow pores appeared to grow along the microrough surface of the implant, with a point of bone-implant contact visible in the PFIB-SEM volume. In both datasets, there is some evidence of bone-implant separation from resin shrinkage and sample preparation but much of the peri-implant space does not contain mineralized bone, evidencing that there is room for further mineral apposition to occur during osseointegration and potential for further improvement of secondary implant stability.

In the 4 wk dataset, some preferential orientation of canaliculi towards the mineralization front is seen (**Figure 3B**) – an observation also associated with *de novo* bone formation. Hasegawa et al. examined canalicular orientation of osteoblasts and osteocytes and found that osteoblast-associated cytoplasmic processes often have a ‘looping’ geometry extending from the secretory side of their membrane, while cell processes associated with osteocytes extend perpendicularly outward from the cell body to connect with osteoblasts at the mineralization front ^[37]^. In addition, osteocytes adjacent to implant surfaces with microrough or minimally rough surfaces can extend cell processes perpendicularly towards the implant interface, eventually attaching to the implant itself ^[38]^. The directionality of the LCN observed in the implant retrieved after 4 wk indicates that osteocytes near the boundary of the mineralized tissue are presumably maintaining a connection with osteoblasts or other bone cells outside of the mineralized matrix. These osteoblasts would, in theory, lie in the space between the mineralized tissue and the titanium. Alternatively, this space may already be filled with osteoid and resident cells. It is most probably that both distance and contact osteogenesis have been simultaneously captured within this FIB-SEM volume due to the intimate contact of bone and titanium in the upper quadrant of this dataset, and large gap between bone mineral and implant surface elsewhere.

After 12 wk of implantation (**Figure 3C** and **D**), the lacunocanalicular network is more densely distributed through the bone matrix, occupying 6.8 vol% of the bone in the 12 wk tomography dataset compared to 3.7% in the 4 wk dataset. Baseline lacunocanalicular porosity levels in mature human femoral bone have been measured at 1.45 vol% and 2.05% utilizing a technique with similar voxel size in synchrotron computed tomography ^[39,40]^, which is a value considerably lower than what is present in newly mineralizing tissue at the implant interface in these PFIB-SEM datasets. A net increase in lacunocanalicular volume density at both 4 wk and 12 wk is expected based on prior work from Shah et al. ^[41]^, who reported high osteocyte counts adjacent to near-cortical implant interfaces in porous titanium (compared to similar locations in solid titanium and within the native bone). For each osteocyte, Shah et al. ^[41]^ also reported a higher number of canaliculi per osteocyte at the implant interface than in the native bone, characterizing tissue with high LCN fraction at the two bone-implant interfaces as newly formed rather than a fragment of pre-existing bone. Beyond the boundary of the mineralized tissue, there also appears to be a number of cells with phenotypes in between that of osteoblasts and osteocytes visible using backscatter imaging without application of a staining protocol (**Figure S3** and **Supporting Information**). One important distinction is that the respective ages of 4 wk and 12 wk correspond to the age of the bone defect and the time of implantation rather than the precise age of the bone tissue itself. While the implant was placed in the tibia for either 4 or 12 wk before retrieval, it is certainly possible that the regenerated bone is less aged than 4 or 12 wk.

Two unique topographies at the boundary of the newly formed bone are seen in the 3D view of the bone-implant interface after 4 wk of implantation (**Figure 4A**-**B** and **Video S1**). Subregions of the mineralizing bone tissue are shown using the 3D renderings in **Figure 4C** and **4D**. One such subregion presents with a rough and granular morphology in 3D. The neighbouring subregion presents with an overall smoother morphology, with only small, elongated protrusions from the mineralized bone. The rough morphology is typical of actively mineralizing bone surfaces, where quasi-spherical calcium phosphate precipitates are observed in the form of calcospherites/calcospherulites ^[42]^, having a mean diameter of roughly 420 nm ^[43]^. Isolation of mineralizing features using connected-component analysis in the implant retrieved after four weeks is able to identify individual mineral ellipsoids or aggregates of small ellipsoids that are close to, but not yet attached to the mineralized bone matrix. 3D rendering of select clusters (**Figure 4E**) shows a wide range of heterogeneity with respect to their size and shape. Newly evolving mineral sometimes appears to present in the ellipsoidal shape described in mature bone tissue ^[4]^, but other times presents in abnormal clusters in conditions such as hypophosphatemic bone ^[5]^. Considering the leporine bone here is otherwise healthy apart from the bone defect, it appears as though these mineral ellipsoids originate from an inconsistent architecture in newly formed tissue **(Figure S4)**. These clusters have a wide distribution with respect to their 3D aspect ratio, supporting what Shah et al. presents in 2D relating to the lack of growth in any preferential direction during ongoing bone mineralization ^[44]^. It is important to note that these isolated clusters perhaps can undergo further growth or aggregation before connecting to the bulk of the bone tissue and that these morphometric parameters are not necessarily representative of mature mineral clusters within the bulk of the mineralized matrix.

**Figure 4.**
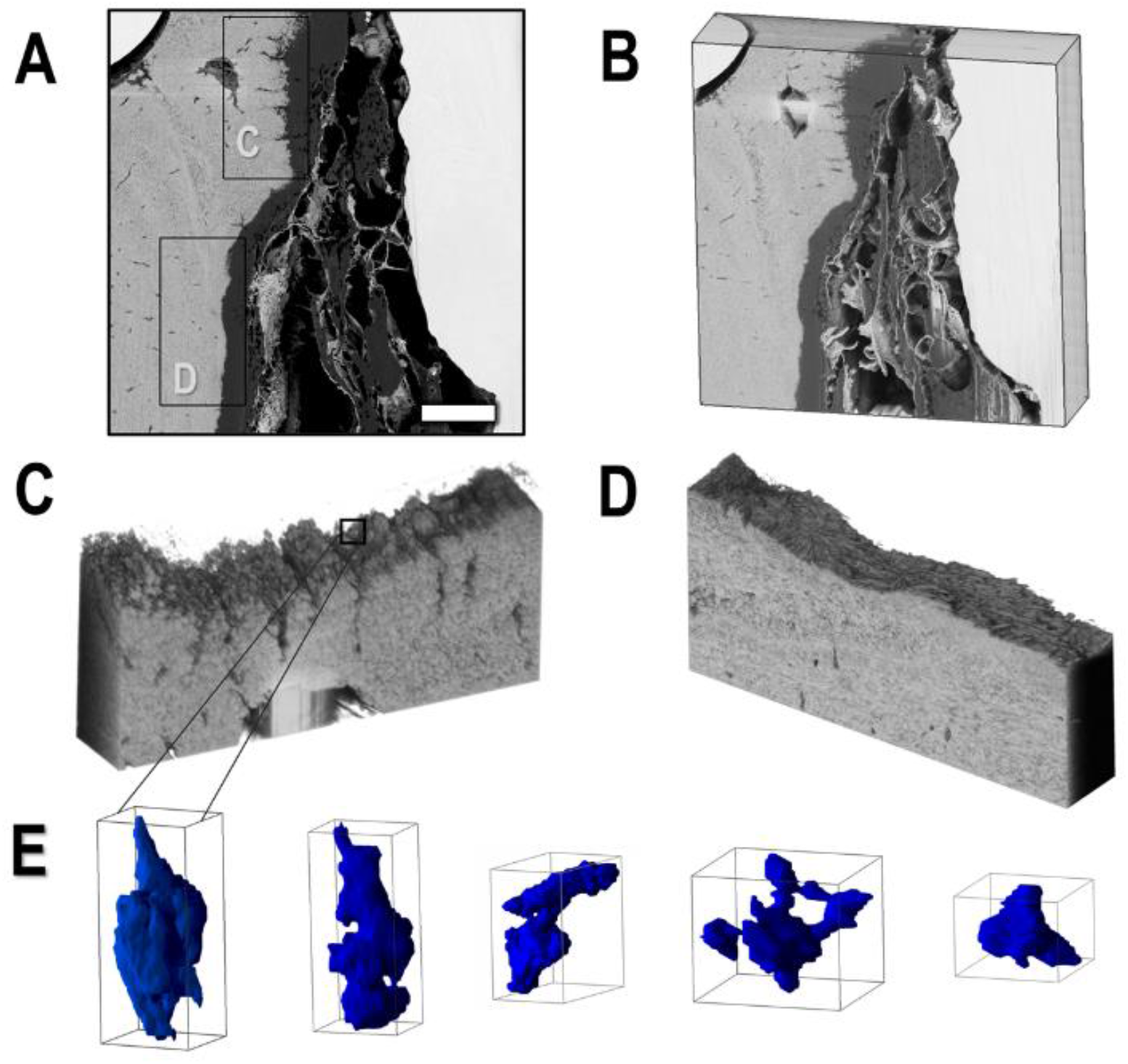
Mineral evolution and topographies at the boundary of the mineralized tissue after 4 wk. (A) Two neighbouring regions at the periphery of the mineralized bone matrix with differing degrees of surface topography. (B) 3D view of the bone-implant interface. (C) 3D topography of the granular or actively mineralizing surface adjacent to the implant interface, with distinct shapes attributed to evolving mineral clusters or foci. (D) Smoother 3D topography, with flatter and elongated protrusions, at a dormant mineralized surface near the implant interface. (E) 3D reconstruction of select mineral clusters at the active site of mineralization, detailing their heterogeneity in size and shape.

The second subregion at the boundary of the mineralized bone, with the smoother topography and more mesh-like organization of mineral, is also characteristic of a mineralizing surface. Examination of sutures in young murine calvaria showed three stages of mineral shape evolution at these smoother mineralized surface in developing bone ^[45]^, presenting with either *(i)* repeating ellipsoid units; *(ii)* a continuous interwoven mesh of mineralized matrix; or *(iii)* some intermediate combination of mesh and ellipsoids. The mesh-like structure in **Figure 4D** can be associated with earlier mineralization events, such as in the floor of osteocyte lacunae ^[45]^, meaning that the mineralizing boundary in **Figure 4D** is perhaps associated with a more dormant mineralization front near the titanium implant due to its intermediate combination of protruding features and meshwork structure.

### 3.3 Mineral Ellipsoid Organization at the Bone-Implant Interface

As noted above, HAADF-STEM imaging of the FIB lift-out near the bone-implant interface has revealed mineral dense domains resembling ellipsoidal structures. Similar mineral ellipsoids are well documented in normal/healthy and diseased bone tissue ^[3–5,46]^. According to the literature, the 2D presentation of a singular mineral ellipsoid depends on its sectioning orientation through the cluster. Longitudinal sections through an idealized cluster will produce the ‘marquise’ or elliptical shape shown in **Figure 5A**, oblique sectioning orientation will produce ellipses with skewed proportions depending on the sectioning angle (**Figure 5B**), while transverse sections will appear as circular (**Figure 5C**), and it is frequently described as the “rosettes” or “lacey” motif ^[47] [48]^. In general, the tip-to-tail orientation (along the long axis) of these mineralized clusters runs parallel to the collagen fibril bundles within the same domain to create the in-plane and out-of-plane motifs. As shown in **Figure 6B and 6C**, some of these distinct mineral ellipsoid motifs were observed near the implant interface.

**Figure 5.**
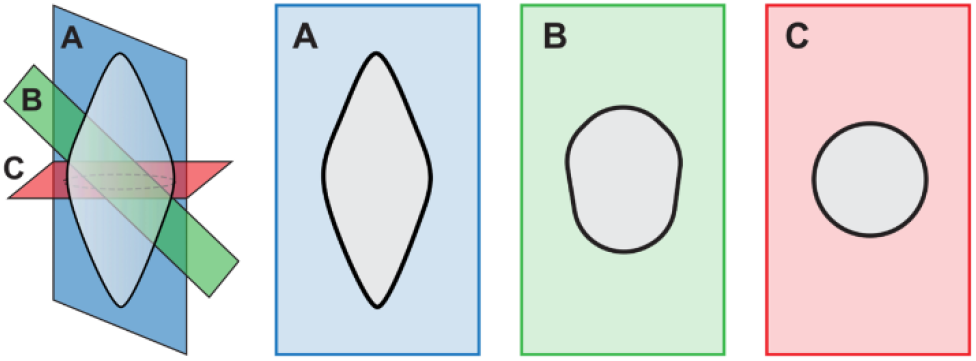
Sectioning planes in a 3D mineral ellipsoid. Sectioning planes of varying angular orientation in a mineral ellipsoid show different ellipsoidal motifs depending on the viewing angle. The elliptical motif in (A) and rosette motif in (C) are parallel and orthogonal to the long axis of the ellipsoid, respectively. Oblique sectioning planes tend to create oblong projections as in (B).

**Figure 6.**
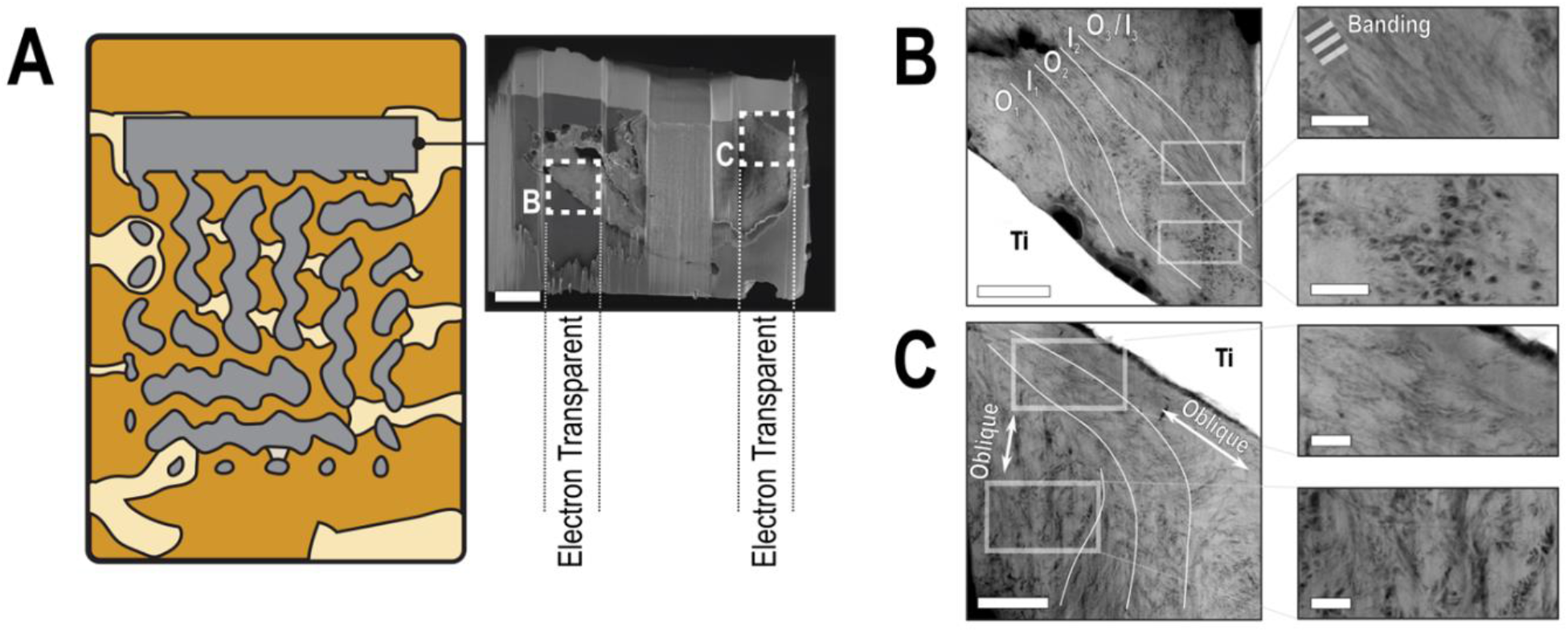
2D mineral twisting motifs at the implant interface after 12 wk. (A) Specimen orientation and sampling location for FIB lift-out. Two electron transparent windows allow for HAADF-STEM imaging. (B) HAADF-STEM imaging shows direct bone-to-implant contact with an alternating array of mineral ellipsoid orientation. Short-order lamellae with unidirectional in-plane collagen orientation (I) and out-of-plane collagen orientation (O) are visible adjacent to the implant. Collagen banding is visible in the inset where collagen lies within the imaging plane. (C) HAADF-STEM imaging of the bone-implant interface in the second electron transparent window. Mineral ellipsoids are obliquely oriented through most of this region. Mineral clusters near the titanium also pack with their long axis parallel to the implant interface. Fewer lamellae are visible in this region, thereby indicating fewer collagen orientation shifts before achieving a unidirectional ellipsoid orientation as distance from the implant interface increases.

Further investigation of the 12 wk sample through HAADF-STEM imaging shows that the mineral ellipsoids were packed into layers of alternating orientation with a thickness of roughly 1-2 μm (**Figure 6**). The layer closest to the implant has a dominant out-of-plane collagen fibril orientation, denoted as O_*1*_ and presenting with a lack of collagen banding, with the long axes of ellipsoids also extending out of the imaging plane. An adjacent region with seemingly similar collagen orientation, based on ellipsoid orientation rather than collagen banding, is shown in the second inset of **Figure 6B**. The intervening layer of mineral ellipsoids, highlighted in the first inset of **Figure 6B**, is instead encapsulated within a more in-plane collagen fibrillar matrix (I_*1*_), where collagen banding is visible due to gap and overlap zones in the collagen ^[49]^. Lamellar packing of these alternating layers occurs in the next layers of mineral for a final structure of *O_1_* → *I_1_* → *O_2_* → *I_2_* within the first 2 μm of the titanium interface. Within each of these first few layers, the collagen fibrils are believed to lie at relatively small angle to the implant interface in either of the in-plane and out-of-plane orientations. Beyond 2 μm, however, a new motif (O_3_/*I_3_*) extends into the bulk of the tissue where the long axes of the mineral ellipsoids (and collagen fibril bundles) no longer lie parallel to the titanium surface and are instead oriented in a more oblique fashion. The introduction of a titanium implant appears to perturb the natural co-aligned and long-range ellipsoidal packing order present in mature bone tissue, with clear demarcations between zones of similar ellipsoidal orientation near the implant interface and a twisting motif between layers.

Within the first 2-3 μm of the implant interface in the second electron transparent window (**Figure 6C**), collagen fibrils still lie approximately parallel to the implant based on the major axis of the oblong mineral ellipsoids (first inset of **Figure 6C**). However, beyond this first zone of co-oriented mineral ellipsoids, there are no subsequent lamellae where collagen fibrils lie parallel to the implant interface. Instead, an abrupt shift occurs in collagen fibril orientation where collagen becomes rather disordered, sometimes orienting the ellipsoid major axis towards the implant interface (second inset of **Figure 6C**). This orientation extends into the tissue with longer-range order than that of the alternating collagen/mineral orientation in the near-implant zone. Short-order periodicity, with corresponding mineral ellipsoid orientation shifts, has been observed in mineralization fronts by Buss et al. ^[5]^, but here we observed a similar phenomenon occurring at the implant interface, albeit not universally. Two-dimensional imaging with HAADF-STEM provides a snapshot of any order and disorder in collagen-mineral packing arrangements ^[50]^ but heterogeneity in structural elements is often better visualized in 3D using techniques such as PFIB-SEM tomography.

A 3D reconstruction of mineralizing bone tissue at the interface of the implant after four weeks of implantation is shown in **Figure 7** and **Video S4**. Removing the titanium region reveals the underlying orientation of mineral ellipsoids directly at the implant interface (**Figure 7B-C**) and slightly further into the tissue (**Figure 7D-E**). As illustrated in **Figure 7C**, there is a clear demarcation between two neighbouring arrays of mineral ellipsoids at the implant interface, each with their own unique orientation and some misorientation angle between arrays. However, examination of 3D ellipsoids further into the tissue shows that the misorientation between these two neighbouring arrays is quickly resolved into that of a unidirectional ellipsoid orientation.

**Figure 7.**
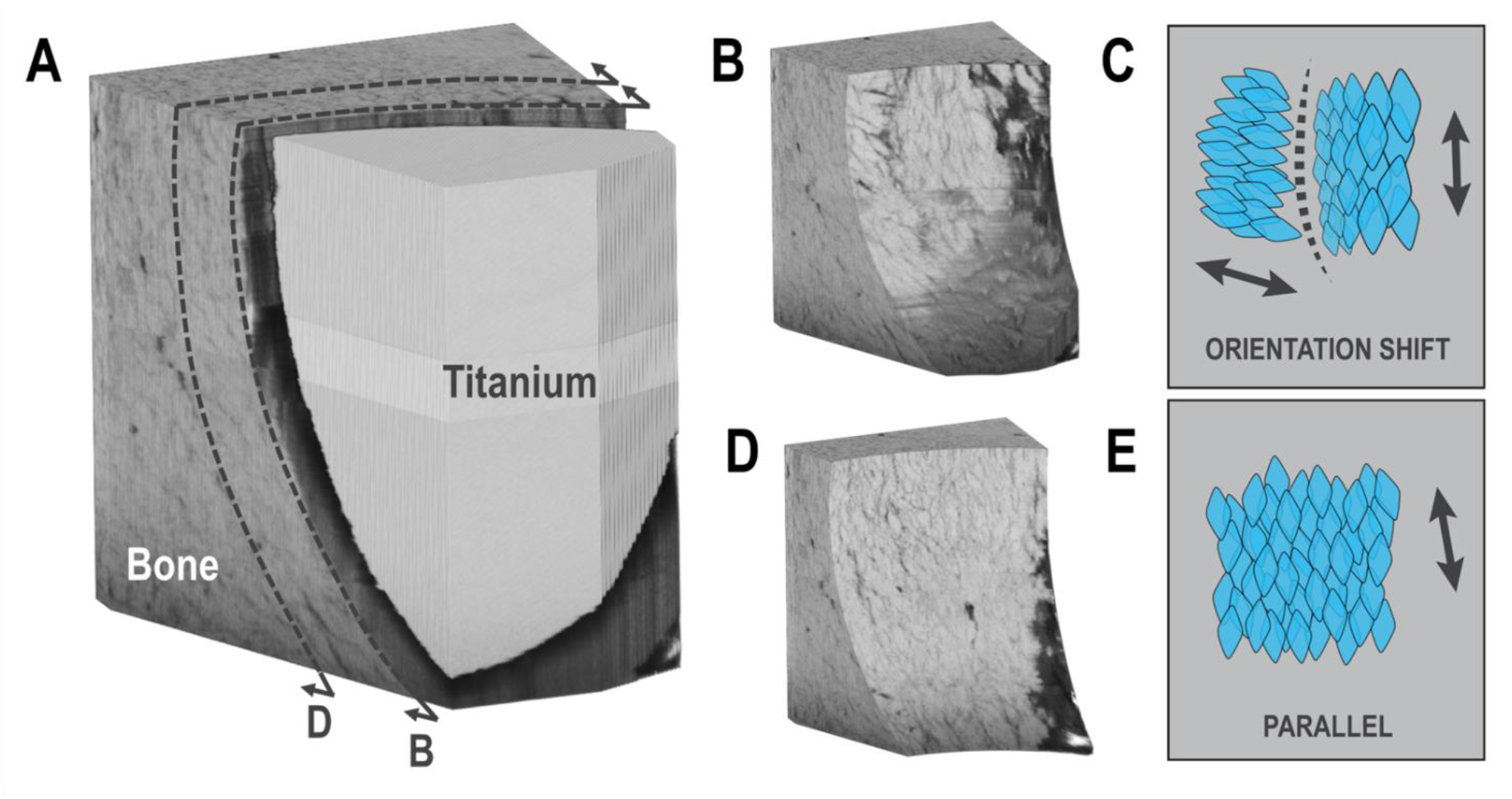
3D mineral twisting motifs at the implant interface after 4 wk. (A) 3D view of bone appearing at the titanium interface from PFIB-SEM with indicated clip planes. (B-C) Examination of mineral nearest the implant interface shows two separate packing arrangements of mineral ellipsoids. An abrupt mineral-deficient region delineates the boundary between these packets of mineral ellipsoids. A size gradient exists across the rightmost array of ellipsoids. (D-E) Slightly further into the tissue, this misorientation angle between ellipsoids is resolved. Further from the implant, the two distinct arrays of ellipsoids merge to form a single array with mostly parallel mineral ellipsoid orientation.

Despite this, it is important to note that each of these two adjacent packing arrangements still have collagenous arrangement that lies parallel to the implant interface based on the major axis of mineral ellipsoids. Also of note is that ellipsoids within a single co-aligned array appear to decrease in size as they approach the neighbouring array, and that this intersection between arrays (visible in **Figure 7B-C**) contains a higher content of organics or pore space than within the bulk of either ellipsoid array.

Near naturally-occurring biological interfaces, namely at the boundary of osteonal lamellae, collagen orientation shifts in the manner of twisted plywood ^[51]^ or Bouligand structure^[52]^ – which is known to add toughness to composite materials ^[53]^. In the bone retrieved from leporine tibiae, we observe a similar twisting effect based on ellipsoid orientation changes that can occur near the interface of the titanium implant. Within the context of the alternating collagen orientation and ellipsoid packing observed in both the PFIB-SEM and HAADF-STEM, these shifts in mineral ellipsoid orientation may provide some form of mechanical advantage and improved mechanical properties of the early bone during osseointegration. Here, we speculate that the collagen and ellipsoid twisting motifs near the implant interface may improve the mechanical properties of the most interfacial bone tissue – especially in comparison to a form of interfacial woven bone – thereby preventing implant instability by increasing the energy required for microcrack propagation in the peri-prosthetic tissue during early osseointegration. Indeed, the mechanical properties of bone are known to vary depending on this collagen orientation - where osteonal bone containing collagen orientation shifts of roughly 90° have higher compressive strength than regions containing solely longitudinal fibres ^[54]^. Where peri-prosthetic fracture is one common means of implant failure, the presence of a stable and mechanically mature bone near the implant interface is one important factor and should complement direct bone-to-implant contact in ensuring successful osseointegration of the implant. Within the context of implant design, one important question for future research is to examine how implant surfaces can be modified or optimized to encourage the rapid formation of these twisting Bouligand structures at the ellipsoidal level in interfacial bone tissue.

Understanding the packing orientation of mineral ellipsoids further from the implant interface is much more convoluted in nature. **Figure 8A** shows the distribution of mineral ellipsoids near a band of high-contrast tissue with less clear demarcations between mineral ellipsoids. Adjacent to this tightly packed region, there appears to be aggregates of low-contrast organic or void space between small mineral ellipsoids. Within the zone of the tightly packed ellipsoids, there does not appear to be any of the three defined ellipsoidal motifs presented in **Figure 5** due to the lack of organic material delineating their boundary. Elsewhere in the tissue at 4 wk (**Figure 8B**), the mineral is distributed in small clusters with self-organized regions containing either the rosette or marquise motifs in the imaging plane. After 12 wk of implantation (**Figure 8C-F**), there is much more size variance with respect to the mineral ellipsoid distribution. In **Figure 8C**, a size gradient exists with respect to these ellipsoids in the rosette motif. Large and seemingly disordered aggregates of organic material or void space (**Figure 8C and 8D**) separate the space between large ellipsoids and almost become indistinguishable from nearby canaliculi. Canaliculi have been shown to propagate mainly through confined regions of disordered collagen ^[55]^, but there has been little discussion of whether canaliculi form and propagate through regions of loosely packed mineral clusters as well. As the distance from the region of large organic aggregates increases, the mineral ellipsoids shrink in size until ellipsoid motifs and their boundaries are difficult to discern in this form of large-volume imaging.

**Figure 8.**
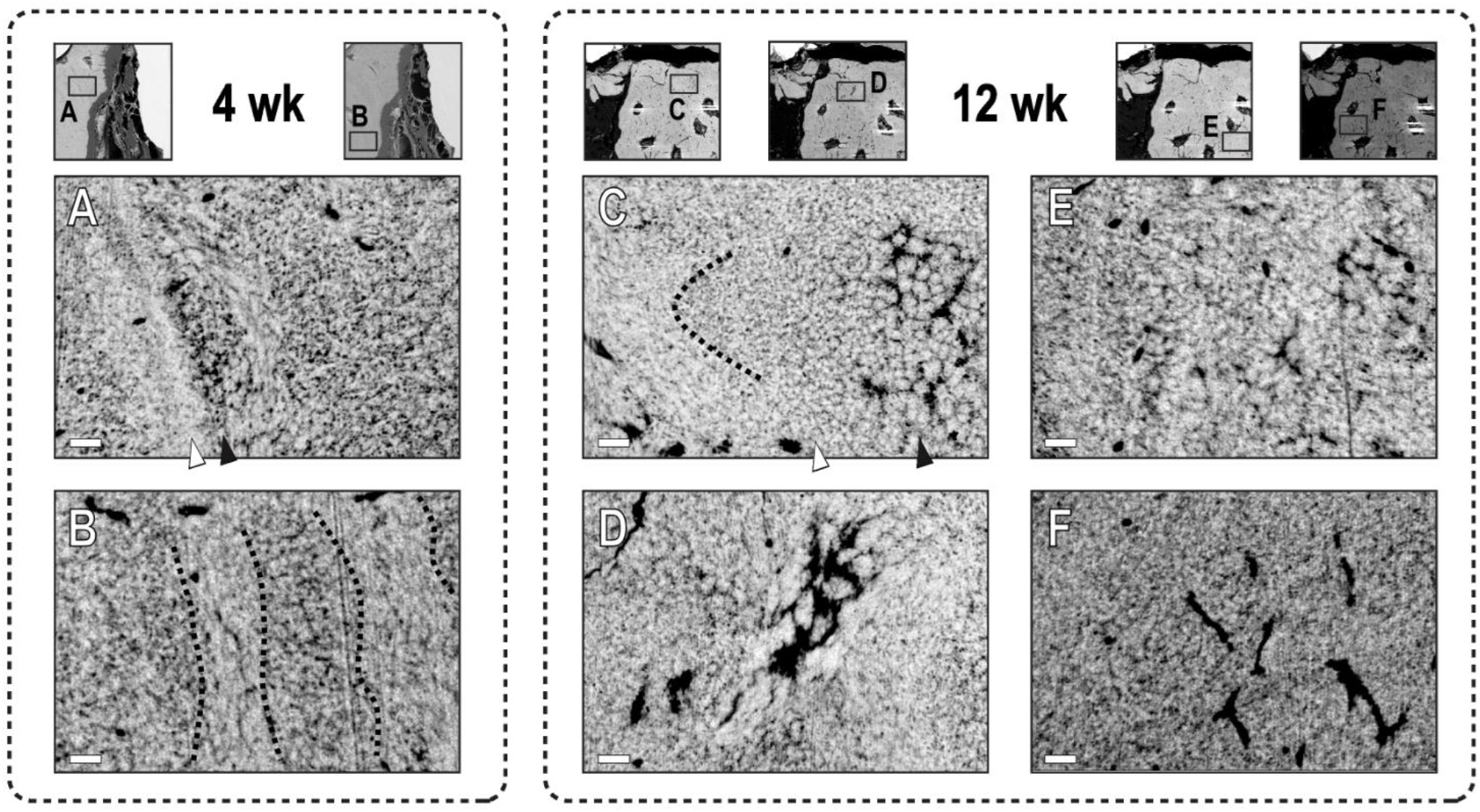
Ellipsoidal heterogeneity is abundant throughout newly formed bone tissue. (A) Large clusters of void space or organic components (black arrowhead) are present adjacent to a band of densely packed ellipsoids (white arrowhead) at 4 wk, disrupting the natural elliptical or rosette patterns on either side. (B) Variation in ellipsoid orientation and organic/void distribution in the bulk of the tissue at 4 wk, separated into regions with common ellipsoid motifs. (C) The size of mineral rosettes varies widely across only a few microns and is small near where the collagen direction shifts after 12 wk. Regions of large ellipsoids are indicated with a black arrowhead and small ellipsoids are denoted with a white arrowhead. (D) Disorganized ellipsoids residing within a largely disordered organic matrix at 12 wk. (E-F) Further evidence of variance in size and organic content in mineral ellipsoids within bone forming near the implant interface. Large aggregates of organic material become indistinguishable with canaliculi in some regions. Scale bars 1 μm.

As with most high-resolution investigations, this study involved a limited number of samples and only selected regions of interest which are assumed to be representative of the remainder of the sample. Despite these limitations, the results of the present work were consistent across all tissue volumes that were examined, and the data showed clear presence of mineral ellipsoids at the bone-implant interface. Nonetheless, future studies involving larger and more diverse sample groups of different age and anatomical locations should be conducted to verify the findings reported here. Additionally, the question of artifacts may be raised by either sample preparation or the FIB liftout and PFIB-SEM imaging techniques. Given the mineral ellipsoidal structures presented here are comparable to those reported in a multitude of healthy and diseased bone tissue by other groups, the artifactual concerns should be diminished. Finally, the lack of targeted staining protocols (heavy-metal staining or others) in the samples makes it difficult to draw concrete conclusions regarding the cellular activity at the bone-implant interface. More studies utilizing other techniques, such as immunocytochemistry, may definitely determine the bone forming process at the interface of these porous materials.

## 4 Conclusions

High-resolution 3D imaging, particularly with PFIB-SEM, affords the opportunity to image multiscale features spanning the nanoscale and microscale continuum. Mineral ellipsoids have recently been characterized in 3D but little is known about their formation and organization at the bone-implant interface. Here, PFIB-SEM tomography and HAADF-STEM are applied to newly formed leporine bone at the interface of a porous titanium implant to visualize mineral ellipsoids with heterogeneous size and orientation during osseointegration. Following retrieval of the porous titanium implants from rabbit tibiae after 12 wk, HAADF-STEM revealed a lamellar packing arrangement of mineral ellipsoids in the first few microns of the implant interface, with alternating orientation between each. A mineral-dense band was also observed directly adjacent to the titanium in some instances. Using PFIB-SEM, a high lacunocanalicular volume fraction was observed in interfacial tissue – a feature characteristic of newly formed bone. Two distinct topographies were observed at active and dormant sites in the boundary of the mineralized tissue, with a wide variety of shapes and sizes in mineral ellipsoids. In the 3D PFIB-SEM datasets, mineral is arranged into ellipsoidal shapes with a heterogenous distribution of both size and shape throughout the newly mineralized tissue. In the bulk of the mineralized bone, this short-order heterogeneity in mineral ellipsoid morphology is also visible, with frequent changes in mineral ellipsoid orientation. These shifts in orientation could possibly be a natural mechanism of increasing fracture toughness to prevent crack propagation and stabilize the implant during osseointegration. Overall, the process of osseointegration occurs as a multiscale 3D process that is best characterized with 3D imaging techniques. Coupling FIB-SEM or PFIB-SEM with sample preparation techniques to highlight non-collagenous protein, organelles, or other organic components features will provide a further understanding of how osseointegration occurs and how mineral is distributed at the interface of both conventional and additively manufactured porous metallic implants.

## Acknowledgements

Work was performed at the Canadian Centre for Electron Microscopy with assistance from Travis Casagrande and Natalie Hamada. The authors would like to acknowledge funding from the Natural Sciences and Engineering Research Council of Canada (RGPIN-2020-05722), Canada Research Chairs Program, Foshan Science and Technology Innovation Project (No. 2018IT100212), and National Natural Science Foundation of China (No. 81801030).

## Notes

### Competing Interest Statement

The authors have declared no competing interest.

