## Supporting Information for "Characterizing Mineral Ellipsoids in New Bone Formation at the Interface of Ti6Al4V Porous Implants"

### Video Captions

***Video S1. Overview of PFIB-SEM tomography at the implant interface after 4 wk of implantation.*** *(i) Slice by slice assembly of the volume with titanium (white), bone (intermediate grayscale), and resin (intervening low-contrast region). (ii) Regions of granular and mesh-like topographies at the boundary of the mineralized tissue. (iii) Heterogeneously sized mineral ellipsoids in the interfacial bone tissue.*

***Video S2. Overview of PFIB-SEM tomography at the implant interface after 12 wk of implantation.*** *(i) Slice by slice assembly of the volume with titanium (white), bone (intermediate grayscale), and resin (low-contrast layer). (ii) Heterogeneously sized mineral ellipsoids in the interfacial bone tissue. (iii) Possible presence of cells beyond the boundary of the mineralized bone tissue – in the absence of heavy-metal staining protocols.*

***Video S3. Region in the PFIB-SEM dataset containing cell-shaped features outside of the mineralized matrix.*** *High-contrast features, including one in the shape of a nucleus, are visible within the boundaries of the membrane.*

***Video S4. Bouligand twisting of mineral ellipsoids directly at the implant interface.*** *(i) Examination of the closest layer of bone mineral to the implant reveals two misoriented arrays of ellipsoids, each with a distinct ellipsoid orientation. (ii) The two distinct arrays merge into a single array with uniform ellipsoid orientation as distance from the implant interface increases.*

### Cells Beyond the Mineralized Matrix

The possible presence of cells outside of the mineralized matrix can also observed in the FIB-SEM volume retrieved after 12 wk (**Figure S3** and **Video S3**), where high-contrast regions are observed in the shape of a membrane. An opposing observation has been previously observed with secondary electron imaging of unstained cells under cryogenic conditions, where lipid-rich regions and the cell membrane appear darker than the cytoplasm rather than brighter [56], but not using backscatter detectors. Where small regions of mineral have been observed as mitochondrial granules or vesicle-bound deposits in the intracellular region of osteoblasts [57,58], and osteoblasts close to the mineralization front are known to be mineral-enriched [59], it is possible that these high-contrast regions in the filtered images are associated with either mineral- or phosphorus-rich regions that are common within plasma membranes or sub-cellular organelles [60]. Image processing operations (**Figure S3A**) may help reveal these cellular features without any form of heavy metal staining during sample preparation. The cavitation in the mineralized tissue surrounding the cell in panel *(i)* of Figure 6C is consistent with that of mineralizing osteocytes [61], Type III osteocytes [62], or other phenotypes in the osteoblast-to-osteocyte transition. The transition of osteoblast to osteocyte involves a shift in the presence of several molecular markers as they become mechanosensory cells encapsulated within the mineralized matrix [61]. The two neighbouring cells in panel *(ii)* are separated by only 1 μm or less in some instances throughout the volume – proximity which is atypical of most osteocyte phenotypes. Instead, their flat morphology, size, and location with respect to mineralized tissue could be attributed to bone lining cells [63] or osteoblasts. Without additional staining via osmium tetroxide [64] or other means, it is difficult to determine whether these are cells embedded within a matrix of secreted osteoid, or if these features simply help demarcate the boundary of mineralized bone tissue.

### *Mineral Cluster Nucleation*

**Figure S4** shows the distribution of cluster volumes, distance from the boundary of the mineralized tissue, and aspect ratio from the isolation of the 3D mineral clusters. The mean cluster volume was measured at 0.0095 $\mu m^3$, or an equivalent spherical diameter of roughly 262 nm, with an average distance of 290 nm from the bulk of the mineralized tissue.

### *Supporting Figures*

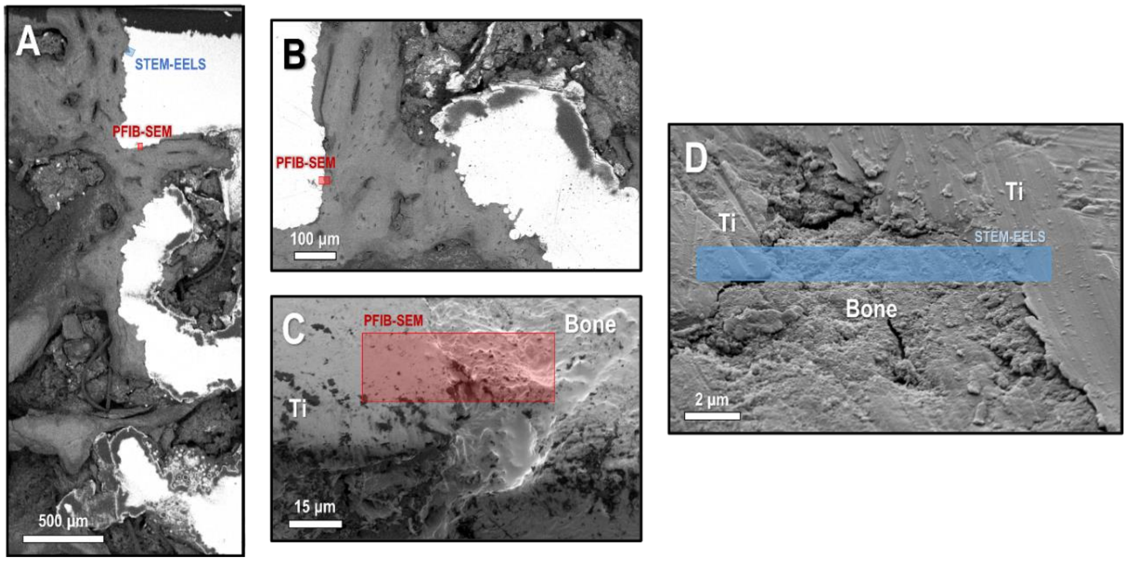


**Figure S1: Location of PFIB-SEM tomography and site of the FIB lift-out for STEM-EELS analysis in the implant retrieved after 12 wk.** (A) Crown of the implant, showing sampling locations for STEM-EELS and PFIB-SEM. (B) Rotated inset showing bone-implant interface sampled for PFIB-SEM. (C) Higher magnification of the PFIB-SEM region. (D) Rotated inset in the crown of the implant showing location of the TEM lift-out.

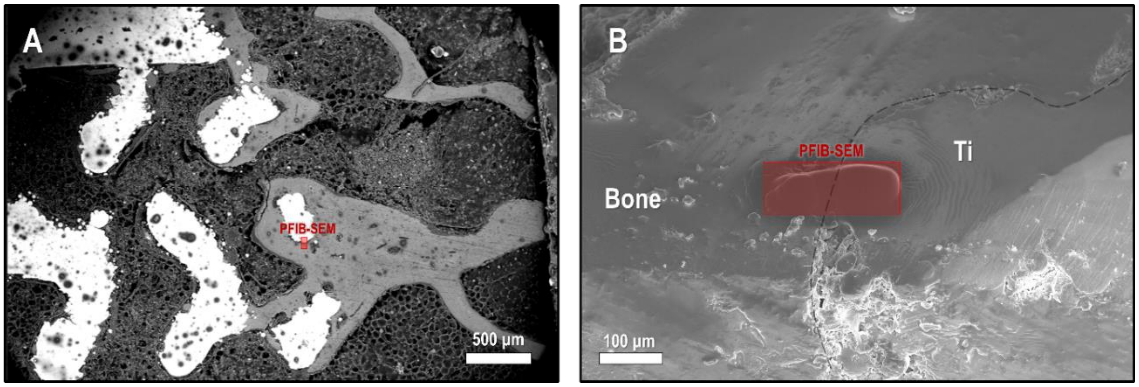


**Figure S2: Location of PFIB-SEM tomography in the implant retrieved after 4 wk.** (A) Broad site overview. (B) Bone-implant interface with deposition of protective carbon layer for PFIB-SEM.

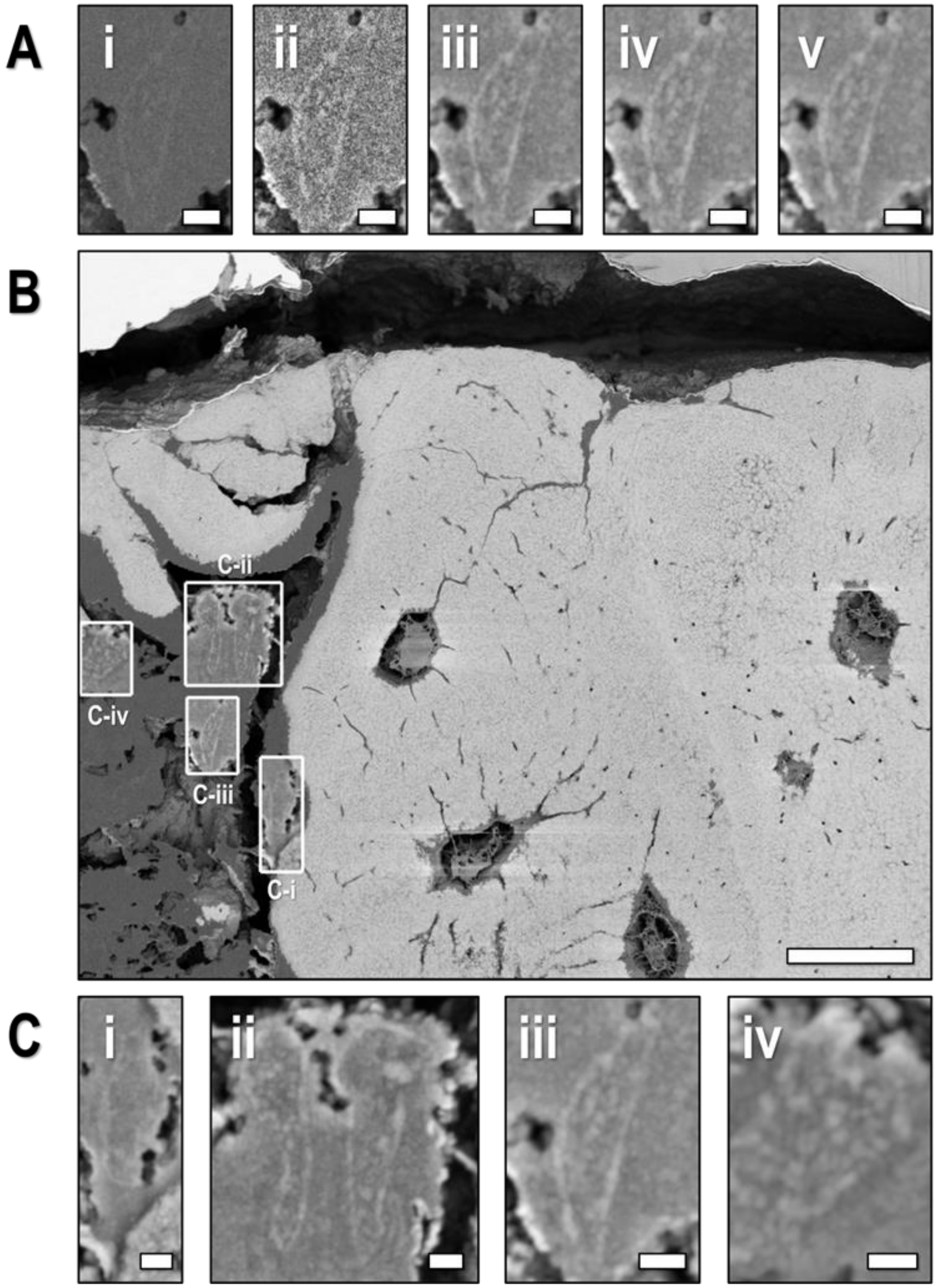


***Figure S3. High-contrast regions in the extracellular matrix beyond the mineralization front.*** *(A) Sequential application of Gaussian smoothing, contrast-limited histogram equalization, Gaussian smoothing, maximum filtering, and Gaussian smoothing to highlight cell membranes beyond the mineralization front. (B) Location of possible cells outside of the mineralized matrix. (C) Insets showing the morphology of nearby cells in the osteoid or extracellular matrix. Scale bars: (A,C) 1 µm (B) 10 µm.*

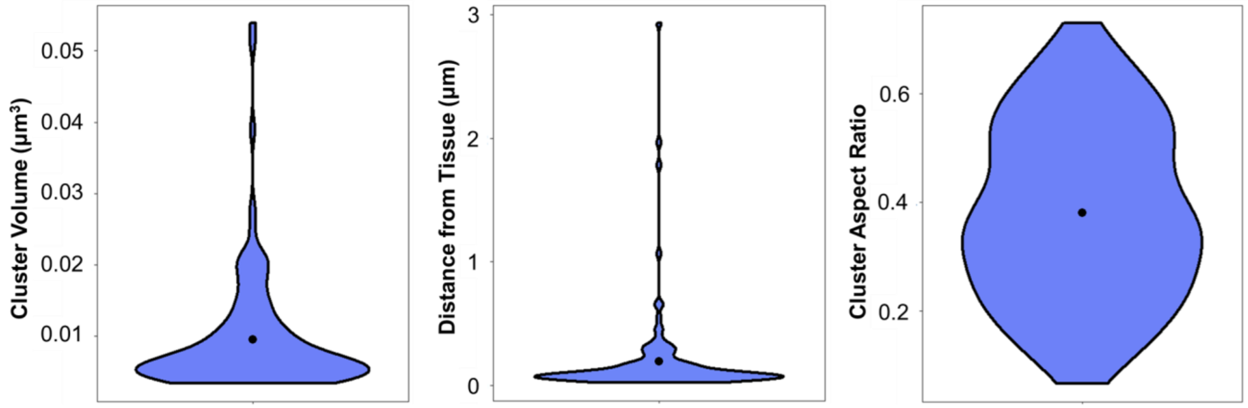


***Figure S4. Mineralization foci morphology.*** *Size, distance to the bulk of mineralized tissue, and aspect ratio of the 3D mineral clusters from PFIB-SEM reconstruction after 4 wk of implantation (n = 116 ellipsoids).*
